# Modeling exercise using optogenetically contractible *Drosophila* larvae

**DOI:** 10.1101/2022.02.16.480715

**Authors:** Arpan C. Ghosh, Yifang Liu, Sudhir Gopal Tattikota, Yanhui Hu, Norbert Perrimon

## Abstract

The pathophysiological effects of a number of metabolic and age-related disorders can be prevented to some extent by exercise and increased physical activity. However, the molecular mechanisms that contribute to the beneficial effects of muscle activity remain poorly explored. Availability of a fast, inexpensive, and genetically tractable model system for muscle activity and exercise will allow the rapid identification and characterization of molecular mechanisms that mediate the beneficial effects of exercise. Here, we report the development and characterization of an optogenetically-inducible muscle contraction (OMC) model in *Drosophila* larvae that we used to study acute exercise-like physiological responses. To characterize muscle-specific transcriptional responses to acute exercise, we performed bulk mRNA-sequencing, revealing striking similarities between acute exercise-induced genes in flies and those previously identified in humans. Our larval muscle contraction model opens a path for rapid identification and characterization of exercise-induced factors.

## Introduction

Exercise and increased physical activity can both prevent and ameliorate the pathophysiological effects of a number of metabolic and age-related disorders. In particular, physical inactivity contributes significantly to age related loss in muscle mass and strength, is strongly associated with the onset of type 2 diabetes and cardiovascular disease, and is a reliable predictor of all causes of mortality [1–8]. Not surprisingly, increased physical activity and exercise remain some of the most prescribed interventions for fighting metabolic and age-associated diseases [3–5, 9–12]. Increased physical activity can prevent to some extent age-associated loss in muscle strength, reduce mortality rates, improve cardiac function, and improve insulin sensitivity [3–5, 12, 13]. Additionally, exercise is beneficial to various pathologies including inflammatory disorders, certain cancers, and osteoporosis [10, 14–16]. Despite the well documented beneficial effects of increased physical activity, our understanding of the molecular mechanisms that mediate these beneficial effects still remains limited.

Several “omics” approaches have been reported that shed light on the role of muscle as a source of exercise-induced humoral factors called “myokines” or “exerkines” [11, 17, 18]. Myokines, such as irisin [19, 20] and meteorin-like [21], are involved in energy metabolism and browning of white fat. In addition, another myokine, apelin, has been demonstrated to improve muscle function by reversing age-associated sarcopenia [22].These studies underscore the diverse roles of myokines in endocrine, paracrine, and autocrine signaling [23]. Furthermore, responses to endurance and strength-based training involve systematic coordination and communication among multiple organs [24], raising the question whether these events during exercise are brought about by myokines. Interestingly, different exercise modules can induce different myokines [18], suggesting that muscles respond differently depending on the type of exercise. Such observations warrant a deeper investigation into how different myokines exert their beneficial effects on various organs.

To date, a number of vertebrate species including humans have been recruited to characterize circulatory myokines in response to different exercise modules. However, in these studies it is difficult to take a systematic look at every myokine in an *in vivo* setting. The availability of *in vivo* genetic model systems would be extremely valuable to characterize the response of muscles to varying degrees of exercise. *Drosophila* has been shown to be an efficient model system to understand the biology of exercise (see reviews [25–27]). For instance, exercise training in young flies can improve age-associated decline in mobility [28]. Of note, the fly ortholog of the vertebrate exercise response gene *dPGC-1α* / *spargel* (*srl*) has been implicated in the physiological effects of endurance exercise [29], suggesting an evolutionarily conserved mechanism of exercise biology. Interestingly, it has also been shown that activating octopaminergic neurons or feeding flies with octopamine can fully substitute for exercise in sedentary adult flies [30]. Therefore, the use of a genetically tractable model organism such as *Drosophila* may allow fast and highly reproducible functional analyses of various cell types, tissues, and genes encoding evolutionarily conserved myokines *in vivo*. However, most of the exercise studies have been modeled around ‘endurance’-based training in adult *Drosophila.* The physiological effects of strength-based or acute exercise in *Drosophila* adults or larvae have not been well addressed. Importantly, besides addressing the physiological aspects of exercise, a systematic evaluation of the transcriptomic architecture of muscles upon exercise in *Drosophila* has not been reported.

Here, we used *Drosophila* larvae to establish an optogenetically-inducible muscle contraction model with the following advantages: 1) larvae are translucent under white light, such that light can pass through the larval cuticle, allowing easy access to motor neurons present in the central nervous system; 2) the larval cuticle is soft, making it possible to induce contraction of all of the body wall muscles at once without causing injury; and 3) larval hemolymph and tissues are easily accessible for biochemical, cell biological, and omics-based approaches. Exploiting such benefits, we demonstrate the application of optogenetic muscle contractions (OMCs) to *Drosophila* larvae and identified several evolutionarily conserved myokine-encoding genes by RNA-sequencing and the signaling pathways that are active in muscles after acute exercise regimens. Further, we identified striking similarities between exercise-induced genes in flies and in humans. We catalog several acute exercise-induced putative myokine-encoding genes, which may aid in the systematic characterization of their roles in beneficial effects of exercise and in addressing the nature of inter-organ communication networks during exercise.

## MATERIALS AND METHODS

### Fly stocks and husbandry

The *OK6-Gal4*^*bc*^ line was generated by backcrossing the original *OK6-Gal4* (BDSC:64199) line to a *w*^*1118*^ control fly line for 7 generations. The *UAS-CsChrim*^*bc*^ line was generated by backcrossing the original *UAS-CsChrim* (BDSC:55136) to the same *w*^*1118*^ fly line for 7 generations. All stocks were maintained at 25°C and experiments were performed at room temperature unless specified otherwise. The endurance training paradigm was performed at 20°C to prolong the larval stage, allowing for a longer regimen of muscle contractions. Egg laying cages were set up by crossing ~150 virgin females and ~120 males in 100 ml volume cages. Crosses were maintained on grape juice plates supplemented with yeast paste and were kept at 25°C under 60% humidity. Egg collections were started from day 4 onwards post mating.

Collections were done for a 3-hour time period every day with a small dab of yeast paste on the grape juice plates. The dab of yeast paste was removed after egg collection and the plates were incubated upside down at 25°C until larvae started hatching. First instar larvae were collected 24 hours after the end of egg collection and either 60 animals were transferred to standard food vials (for acute exercise training) or 30 animals were transferred to all-trans retinal (ATR) containing food (for endurance training).

### Tissue dissections

#### Body wall muscle

Dissections were done in 9-well depression plates with the wells that were half filled with silicone elastomer to allow positioning of the larvae and fillet using insect pins. Dissections were started by adding 1 ml of relaxing buffer [(RB) 1× PBS, 5 mM MgCl_2_ and 5 mM EGTA] to the well and placing a washed and cleaned 3^rd^ instar larvae in the well. The animal was positioned ventral side up and the head and tail of the animals were pinned using insect pins with a gentle stretch along the body. A pair of fine micro scissors was used to cut along the ventral midline of the larvae. A 1 ml pipette was used to squirt the RB inside the animals until all the muscles were visibly relaxed. 4 additional insect pins were used to open the fillet and lay the larval cuticle flat. Internal organs were carefully removed making sure not to injure the muscle attached to the cuticle. Samples were fixed in 4% paraformaldehyde in RB for 20 mins in the same well while the cuticle was still pinned to the silicone bed. Post fixation, the insect pins were removed, and the fillets were transferred to low adhesion tubes for further processing. For experimental larvae that show a strong contraction under the white light, we followed the following additional steps. First, we used a lower intensity of white light to do the dissections. Second, we waited at least 30 seconds to allow the larvae to relax after the initial strong contraction under the bright white light. Third, after the incision along the ventral midline and once the RB had relaxed all the muscles, we readjusted the insect pins to stretch out any slack that was caused by relaxation of the body wall muscles.

#### Fat body

Dissection was performed in a deep-well dish with the well half filled with silicone elastomer. A 20 μm diameter insect pin that was bent by 90 degrees was inserted in the silicone to create a hook that was used to invert the larvae. 1 ml of RB was added to the well, and stage and size matched subject animals were washed and placed in the well. The posterior end of the animal was cut with a pair of micro scissors and two sharp forceps were used to gently invert the animal on the insect pin keeping all internal organs to one side of the body. Subsequently the gut was removed while making sure other organs were not damaged. Tracheae were gently pulled out by holding the cut end of the primary tracheal tubes and pulling them along the length of the body. The carcass with the fat body still attached was then unmounted from the insect pin and transferred to RB in non-adhesion tubes. Samples were fixed in 4% paraformaldehyde in RB for 20 mins before use for subsequent processing. Final dissection of the fat body was performed on slides prior to mounting.

#### Oenocytes

Sample preparation for oenocytes was performed identically to muscle preparation with the exception that animals were mounted with the dorsal side facing up and the incision was made along the dorsal midline.

### Antibody staining

Third instar muscle fillets were permeabilized in 0.1 % Triton-X100 in PBS (PBT) for 30 mins. Samples were then blocked in PBT + 5% BSA blocking solution (BS) for 2 hours at RT. Subsequently primary antibody (anti-glycogen Mouse anti-IgM: 1/100, anti-ATP5α: 1/500) diluted in (BS) was added to the samples. Samples were then incubated over night at 4°C with gentle rotation. The following day, the samples were washed in PBT 4 times (15 mins each). Samples were then incubated with an appropriate secondary antibody (with 1/500 dilution) in BS for 2 hours at RT. Next, the samples were washed again with PBT 4 times (15 mins each) and mounted in VectaShield with DAPI (VectaShield 1200). Samples were mounted on a slide with the body wall muscles facing a coverslip. The coverslip was bridged to avoid squishing of the sample by sandwiching two stacked pieces of scotch tape (3M) between the coverslip and the slide on both ends of the coverslip. Samples were imaged using a Zeiss LSM780 confocal microscope. Images were acquired at RT.

### BODIPY staining of oenocytes and fat body

Dissected and fixed tissues were permeabilized in 0.1% PBT for 30 mins at RT. Samples were washed three times, 5 mins each, in PBS to remove all Triton-X100. A 1 mg/ml stock of BODIPY in DMSO was diluted at 1/500 in PBS and added to the samples. Staining was performed for 30 mins at RT with constant rotation. The samples were then washed with PBS three more times, 5 mins each, and mounted on slides in VectaShield with DAPI (VectaShield 1200). The coverslip was bridged to avoid squishing of the sample by sandwiching two stacked pieces of scotch tape (3M) between the coverslip and the slide on both ends of the coverslip.

### RNA sequencing and analysis

Total RNA was extracted from larval cuticle preparations using a Qiagen RNAeasy mini prep kit as per the manufacturer’s instructions. RNAseq on mRNA-selected samples were performed by the Sulzberger Columbia Genome Center, Columbia University, using an Illumina HISEQ 2500 instrument. We obtained approximately 30 million single-end reads per biological replicate. The quality of the sequencing reads was checked using the FASTQC package and reads were mapped onto the *Drosophila* genome (release 6) using the STAR aligner package. The read counts were calculated using featureCounts. The subsequent read counts matrix was further processed in R software using the Deseq2 package for normalization and differential gene expression analysis. Principal Component Analysis (PCA) was performed by plotPCA function from Deseq2 package using top variance genes. The heatmap for Figure 3B was generated by R package pheatmap. We used Gprofiler2 and cluster profiler to measure gene ontology enrichment and the resulting illustration was done by emapplot function (Figures 3D, 4B) or by cnetplot function (Figure 4C, S3B and S4) from clusterProfiler package. The annotation of the core components of signaling pathways was obtained from GLAD [31] and enrichment analysis was done using a program written in house based on hypergeometric distribution. The heatmap illustration of pathway enrichment result was done using TM4 software suite (http://mev.tm4.org/). For Figures 5B and S5, we used the gene log2 FoldChange and pathway information as input and plotted the bar plot using ggplot2 package and used coord_polar function to convert the standard bar plot to the circle plot. The raw RNA-seq reads have been deposited in the NCBI Gene Expression Omnibus (GEO) database with the accession number GSE196850.

**Figure 1:**
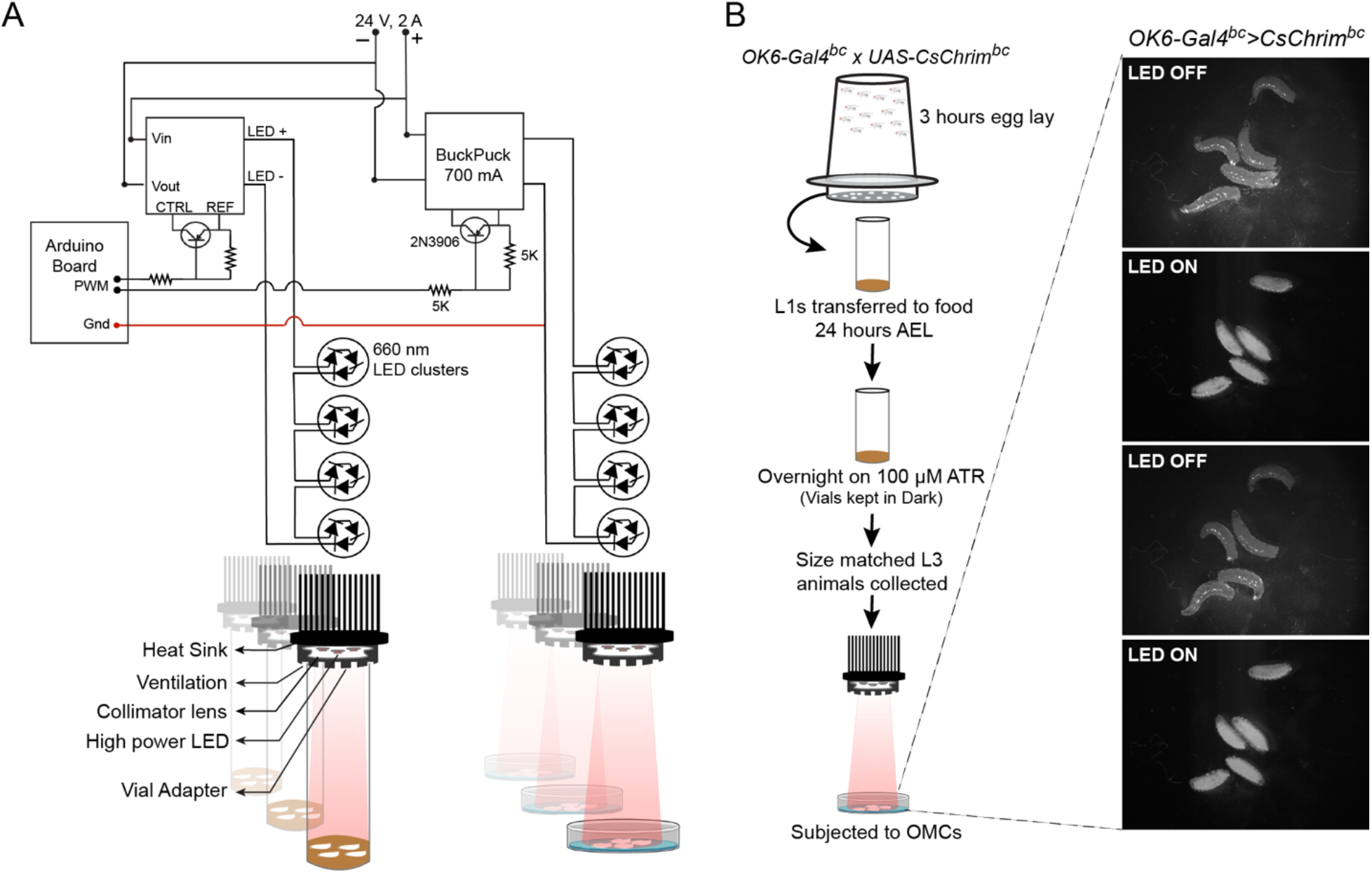
Schematic of the experimental set up for optogenetic muscle contraction (OMC) (A) Schematic of the optogenetic set up used to induce muscle contractions in *Drosophila* larvae. TOP: Circuit diagram of all the components involved in regulating the light emitting diodes (LEDs). The circuit components allow rapid switching of the high-power LEDs using a microprocessor controller such as an Arduino board. (BOTTOM) Schematic of the heat sink, to which the LEDs were attached, and the attached assembly used as a holder for the larval vials. Holders were assembled from laser cut acrylic pieces and are designed to prevent light leak between tubes while allowing air flow to support oxygen supply. (B) Schematic of a basic experimental set up for inducing optogenetic contractions in third instar larvae. The images show snapshots of larvae over-expressing CsChrimson in their motor-neurons (*OK6-Gal4>CsChrim*) when the LEDs were either OFF or ON, demonstrating the synchronized contraction of the larval body wall muscles

**Figure 2:**
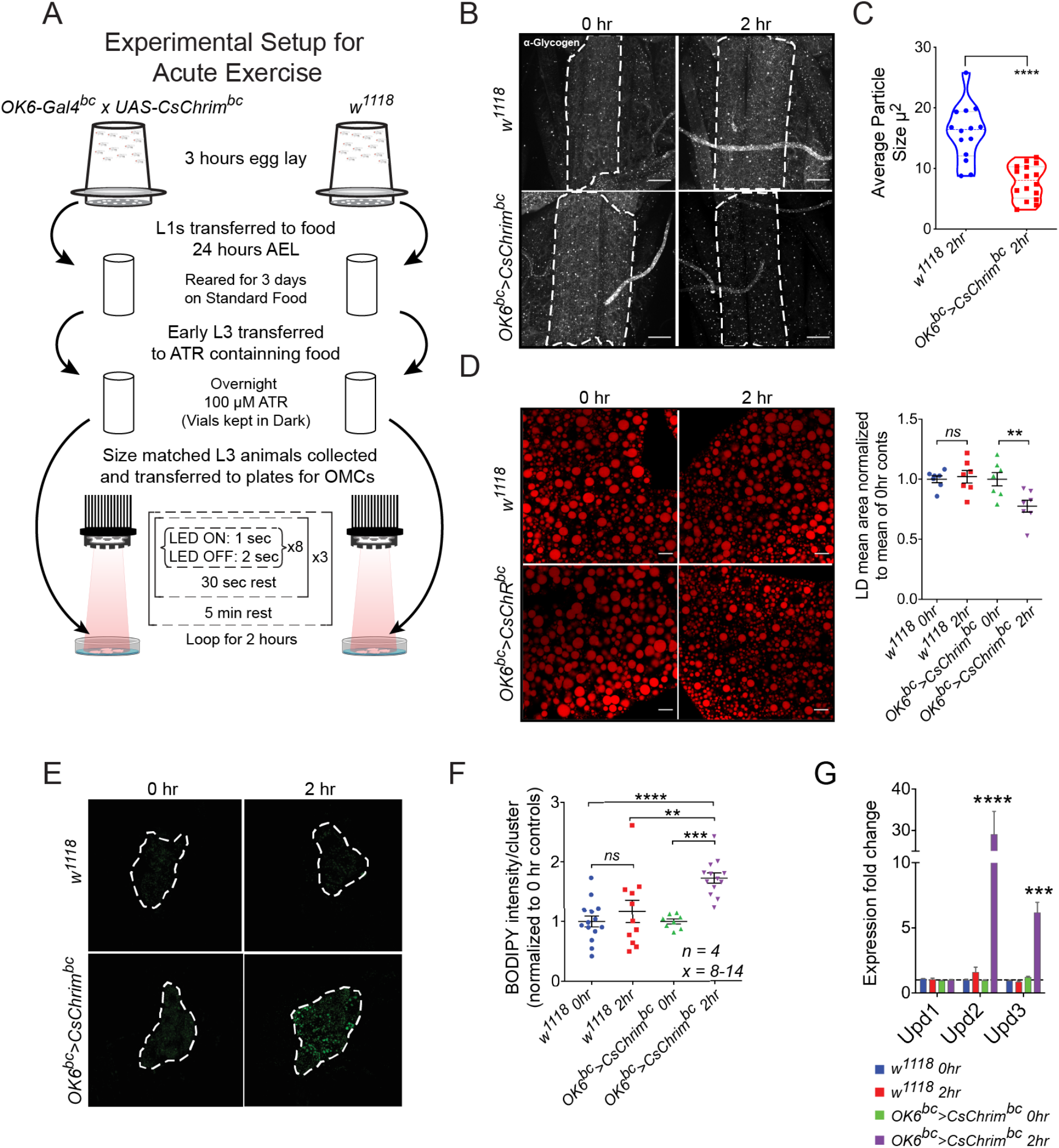
Modeling acute exercise using OMCs. (A) Schematic showing the experimental set up and frequency of OMCs used for modeling of acute exercise in *Drosophila* larvae. (B) Anti-Glycogen antibody staining of larval muscles before and after 2 hours of acute muscle exercise. Control animals do not show a strong change in the levels of glycogen during the 2 hours of light stimulation. Experimental animals however show strong reduction in glycogen staining levels, suggesting a depletion of muscle glycogen reserves. Scale bars = 50 μm. (C) Quantification showing average size of the glycogen particles identified using the anti-glycogen antibody. **** represents a p-value ≤ 0.0001 by Student’s t-test. (D) BODIPY staining of larval fat body cells showing the effect of acute exercise on lipid droplet size. While control animals do not show any change in response to 2 hours of light stimulation, experimental larvae show a significant reduction in the size of lipid droplets in their fat body. The mean area of the lipid droplets at 0 and 2 hours of light stimulation was quantified for both genotypes and is presented in the adjacent graph. N = 7 biological replicates and two-way ANOVA followed by post-hoc test was used for calculating significance. ** denotes p ≤ 0.01. Scale bars = 20 μm. (E) BODIPY staining of larval fat oenocytes showing the effect of acute exercise after 2 hours on lipid accumulation in the oenocytes. (F) Quantification of oenocyte BODIPY staining. Experimental animals show a significant increase in neutral lipid staining in response to 2 hours of light stimulation, but control animals do not. 8-14 oenocyte clusters from 4 independent animals (N=4) were imaged for the analysis. N=4, Two-way ANOVA followed by post-hoc test was used for calculating significance. **** denotes p ≤ 0.0001, *** denotes p ≤ 0.001, ** denotes p ≤ 0.01. (G) Quantification of the mRNA expression levels of *Drosophila* IL6 orthologs, *upd1*, *upd2* and *upd3*, in the body wall muscles of control and experimental animals subjected to 2 hours of light stimulation. *upd2* and *upd3* expression increased significantly only in experimental animals after 2 hours of light stimulation. *upd2* showed ~30 fold increase in expression in response to acute exercise. N=4 biological replicates.

**Figure 3:**
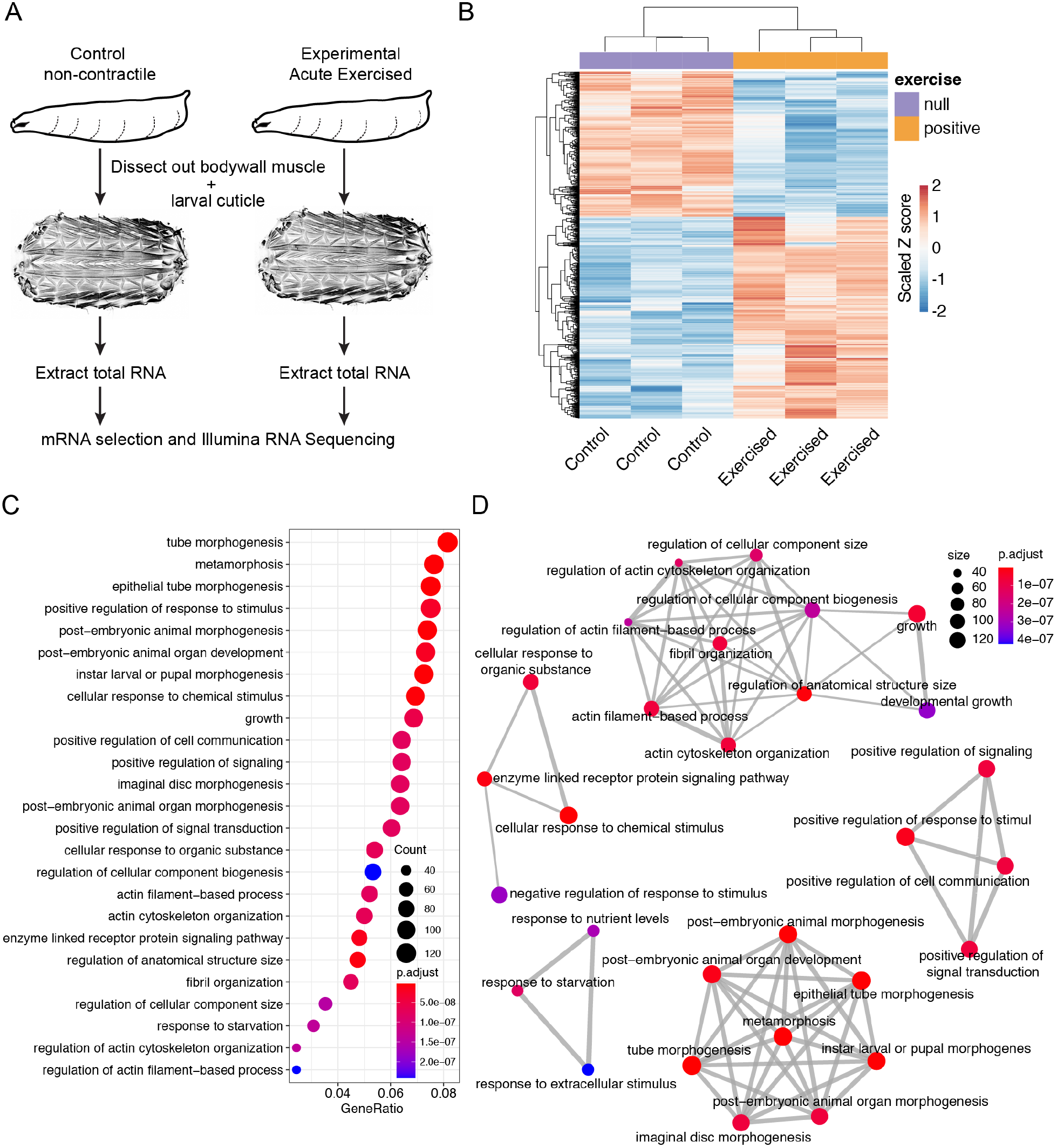
Analysis of all genes for which expression changed in response to acute activity. (A) Schematic of the experimental setup for RNA-Sequencing based assessment of transcriptional changes caused by acute exercise using the OMC model. (B) Heatmap showing all 1190 genes that show a significant change (adjusted p-value ≤ 0.05) of 1.5 fold or more compared to control samples. (C) Dot plot showing the top 25 most significantly enriched GO terms. The color represents the adjusted p-values of the enrichment analysis relative to the other displayed terms (brighter red is more significant) and the size of the terms represent the number of genes that consist the term. (D) An enrichment GO plot showing the relationship between the top 25 most significantly enriched GO terms based on adjusted p-value. Similar terms are grouped together. The color represents the adjusted p-values of the enrichment analysis relative to the other displayed terms (brighter red is more significant) and the size of the terms represent the number of genes that consist the term.

**Figure 4:**
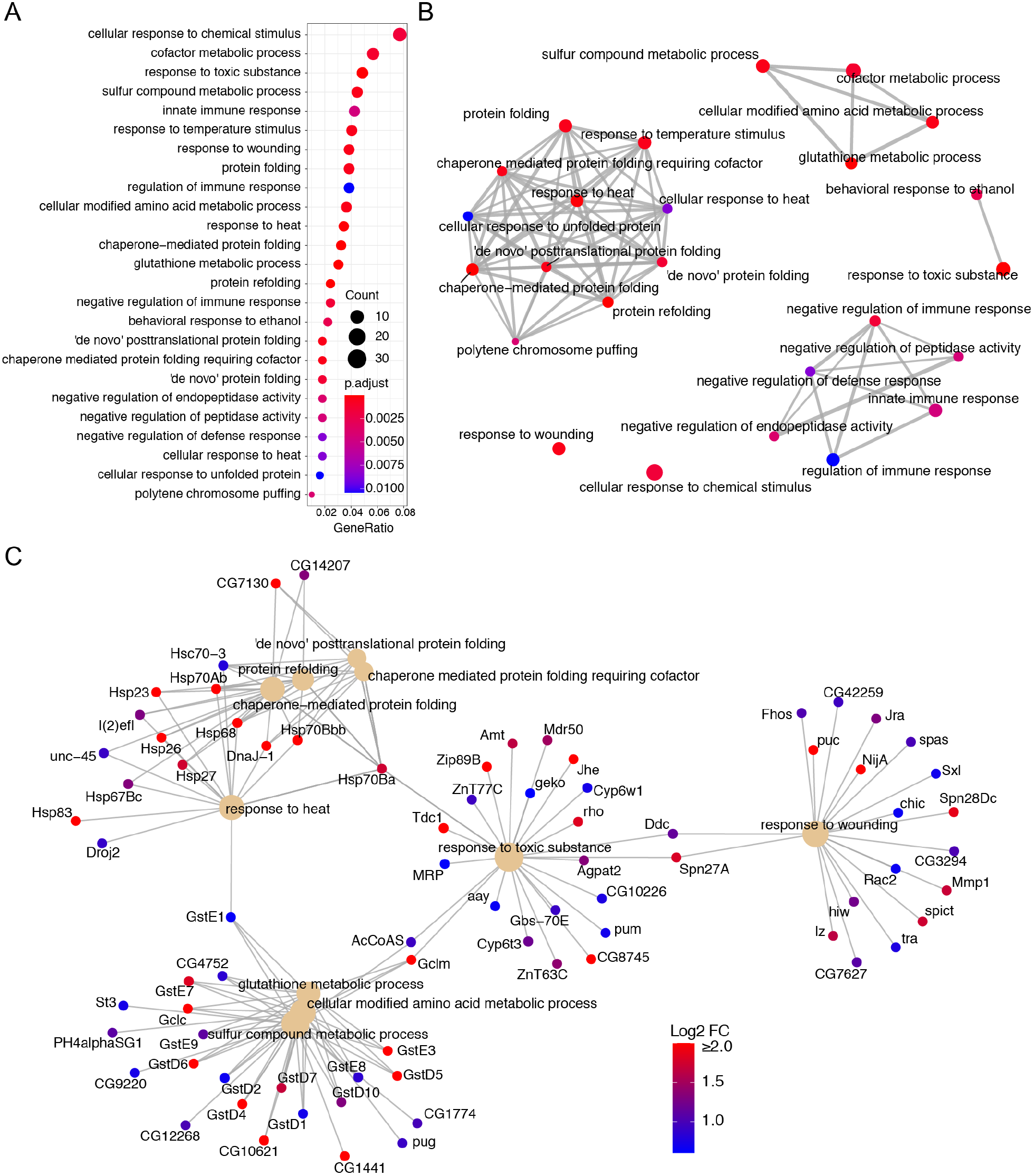
Analysis of genes upregulated in response to acute exercise. (A) Dot plot showing the top 25 most significantly enriched GO terms for genes that show a significant up-regulation of 1.5 fold or more. The color represents the adjusted p-values relative to the other displayed terms (brighter red is more significant) and the size of the terms represent the number of genes that consist the term. (B) An enrichment GO plot showing the relationship between the top 25 most significantly enriched GO terms shown in (A). Similar terms are grouped together. The color represents the adjusted p-values relative to the other displayed terms (brighter red is more significant) and the size of the terms represent the number of genes that consist the term. (C) A cnet-plot showing top 10 GO terms that were significantly enriched for genes that show an upregulation of 1.5 fold or more. The color of gene node represents the log2 value of-fold change with the brightest red representing the log2-FoldChange ≥ 2 and the darkest blue representing a log2-FoldChange of 0.60.

**Figure 5:**
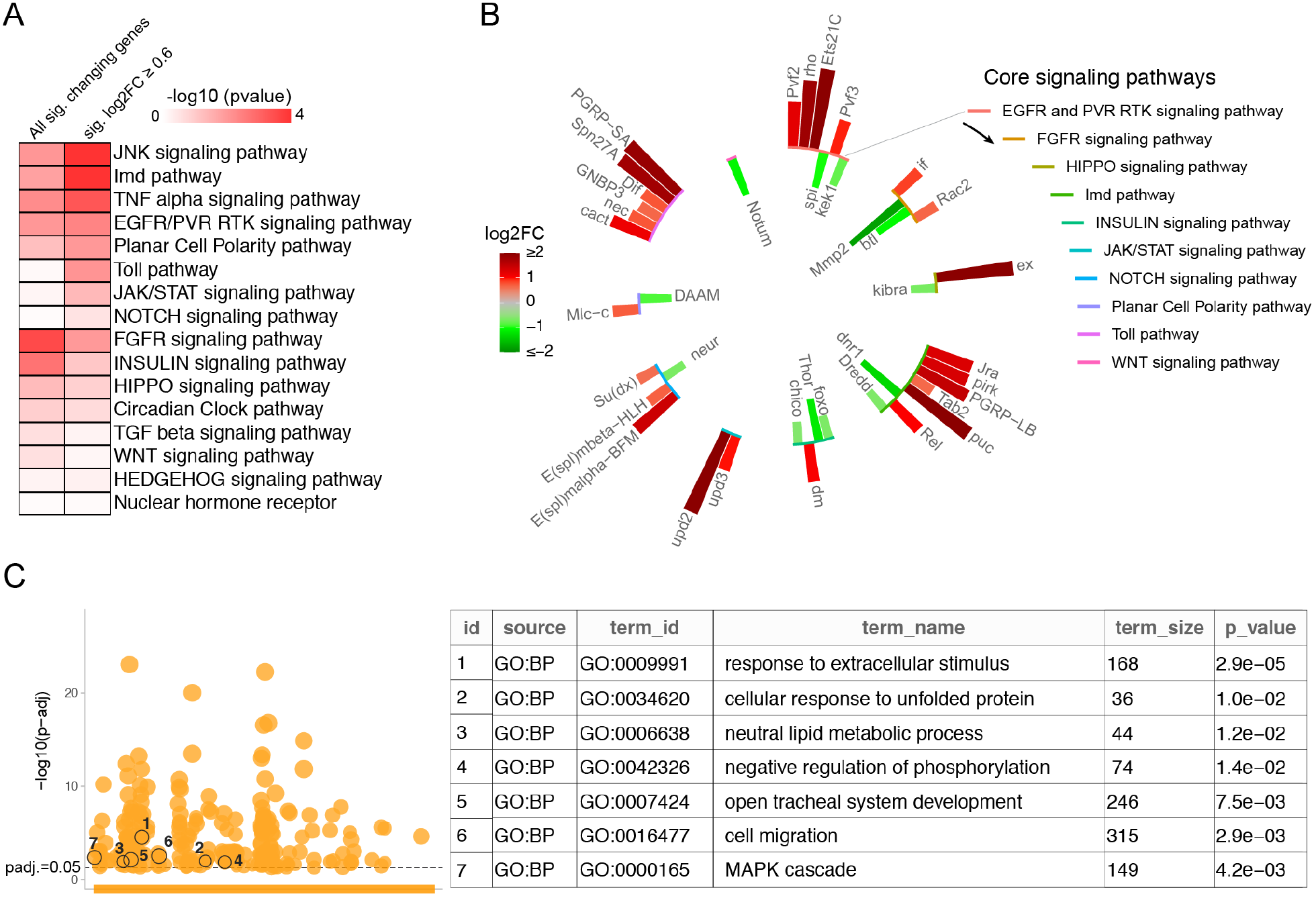
Acute exercise-induced changes in signaling pathways. (A) An enrichment map of core signaling pathways looking for over-representation of component genes in either all significantly (sig.) changing genes or genes that showed a significant and more than 1.5 fold increase in expression. (B) Signaling pathway genes that showed a significant and more than 1.5 fold change in expression. (C) GOSt plot (image on left) showing significantly enriched GO terms related to biological processes with adjusted P values [-log10 (p-adj)] on the y axis. The numbers encircled on the GOSt plot represent the GO terms (see table on the right) that are reported by Pillon et al. (PMID: 31980607) to be commonly enriched amongst genes upregulated in response to aerobic exercise in healthy or metabolically impaired subjects (Humans). These GO terms are also over-represented in our acute exercise induced genes.

## Results and Discussion

### Optogenetic muscle contraction (OMC) model using *Drosophila* larvae

Our setup consists of a series of high-power light emitting diodes (LEDs) that are controlled by an Arduino microprocessor controller via a number of Buckpuck power regulators and switches. Details on the components used to make the OMC setup are available in Table 1 and the circuit diagram for the setup is shown in Figure 1A. Adapter rings that allow attachment of standard narrow *Drosophila* food vials without interrupting air flow were made by mounting laser-cut acrylic pieces around the LEDs. Scaled diagrams for the adapter ring setup are depicted in Figure S1. Optogenetic muscle contractions were performed on animals that were either kept in 30 mm cell culture plates in a thin layer of 2% sucrose or in food vials containing 2-3 mm of food per vial. The adapter ring setup allows attachment of up to 12 vials in a single experiment.

Our standard workflow for an OMC experiment is represented in Figure 1B. Briefly, *Ok6-Gal4*^*bc*^/+; *UAS-CsChrim*^*bc*^/+ (*OK6-Gal4*^*bc*^<*CsChrim*^*bc*^) embryos were collected for a window of 3 hours on grape juice plates. 24 hours after egg lay (AEL), 60 first instar larvae were transferred to standard food vials that were wrapped in aluminum foil to prevent exposure to light and reared at 25°C for 3 days. The evening before induction of OMCs, mid third instar larvae were floated out of the food using 20% glycerol and 30 size matched larvae were transferred to new food vials containing 100 μM all-trans retinal (ATR), a substrate for Channelrhodopsin. Next, the vials were wrapped in foil and the larvae reared at 25°C overnight. The following morning, size matched experimental and control larvae were again selected and placed in 30 mm cell culture plates for inducing OMCs. Representative images of animals that are either relaxed or are contracted in response to 660 nm light stimulation are shown in Figure 1B and Video 1.

### Modeling acute exercise using OMC in *Drosophila* larvae

There are two modalities of physical exercise, endurance training and acute training, which differ in the intensity and duration of the regimens [24]. Endurance training involves low intensity but highly repetitive bouts of training regimens, whereas acute or strength-based exercise typically involves a short duration of high intensity bouts of exercise [32]. Most studies in *Drosophila* focused mainly on endurance-based exercise regimens using adult flies [26] and work addressing acute exercise is relatively scant. Hence, we reasoned that our OMC model could be utilized to mimic acute exercise in larvae, which are ideal for passing light through the translucent cuticle for optogenetics and subsequent induction of body wall muscle contractions. Because acute exercise training is based on the intensity and frequency of exercise bouts, followed by depletion of muscle glycogen [33, 34], we intended to induce acute exercise by inducing rapid muscle contractions and use muscle glycogen as a readout to determine the acute exercise program in larvae. For measuring glycogen levels, we used a previously validated immunofluorescence-based method to visualize muscle glycogen [35, 36]. For re-validating the anti-glycogen antibody in our experimental setup, we first tested the specificity of the antibody after treating the body wall muscles with amyloglycosidase, an enzyme known to deplete glycogen [35]. As expected, anti-glycogen antibody failed to detect any glycogen in the muscles that were treated with amyloglucosidase, suggesting that the antibody-based method to detect glycogen is reliable (Figure S2A-B).

Next, we induced an acute exercise program with a series of repetitive muscle contractions over a period of 2 hours (see schematic in Figure 2A). Briefly, size matched mid-third instar experimental (*OK6-Gal4*^*bc*^>*CsChrim*^*bc*^) and control (*w*^*1118*^) larvae were kept on ATR containing food overnight. On the day of the experiment, the animals were floated out of food using 20% glycerol and transferred to 30 mm petri dishes that contained 2 ml of a 2% sucrose solution in PBS. It is important at this stage to wipe the sides of the plates with a cotton swab dipped in acetone to create a hydrophobic surface that prevented formation of a concave meniscus. The above two conditions prevented the animals from wandering off the plate while making sure that they did not drown in the liquid. The acute exercise paradigm consisted of 8 rapid contractions consisting of 1 sec each of light stimulation interspersed by 2 seconds of no light stimulus followed by a 30 second break (rest). This cycle was repeated 3 times before giving the animals a 5 min of rest period. The entire exercise cycle was repeated in a loop for 2 hours. At the end of the acute exercise paradigm the larvae remained inactive and devoid of any contractile action for a short period of time. The animals were, however, alive as evidenced by a beating heart tube. When transferred to fresh food and dark conditions, the exercised animals recovered and developed to pupae and then adults. Interestingly, adults that emerge from these pupae failed to expand their wings, a phenotype that is very similar to what is seen in flies that are mutant for the genes *bursicon* (*burs*) and *partner of bursicon* (*pburs*) [37, 38]. *Burs* and *pburs* encode cysteine knot proteins that dimerize to form the neurohormone Bursicon that regulates cuticular pigmentation and wing expansion in adult flies. These genes are produced and released from the CCAP neurons in the fly brain and exert their effects on a distinct set of neuronal network that regulate the complex behavior of wing expansion [39]. The *OK6-Gal4* driver used in our experiments is expressed in a subset of the CCAP neurons and the similarity in phenotypes indicate that the activation of these neurons possibly impairs the production of Bursicon or leads to premature release of Bursicon, leaving the adult flies devoid of Bursicon and hence incapable of inflating their wings [40]. This observation highlights one of the potential drawbacks of our OMC model, i.e., that non-specificity of the neuronal driver might cause systemic physiological effects by impacting neuro-endocrine signals.

To determine the physiological impact of our acute exercise paradigm, we monitored muscle glycogen levels in control and exercise animals using the anti-glycogen antibody-based method of detecting glycogen reserves. Relative to control animals, we observed a strong and significant decrease in muscle glycogen levels in exercised animals (Figure 2B-C). Interestingly, when we changed the exercise regimen to endurance-like training over a period of 72 hours (Figure S2C), we observed that the muscles tend to accumulate glycogen levels, in contrast to those of acute exercised animals (Figure S2D, E). This differential depletion or accumulation of glycogen levels is quite similar to that of muscle glycogen levels in humans depending on the exercise regimen [41, 42], suggesting that our larval OMC model is relevant in the context exercise physiology in humans.

The acute exercise regimen also led to a significant reduction in the size of lipid droplets in the larval adipose tissue, indicating lipid mobilization from the lipid stores (Figure 2D). Interestingly, whole animal triacylglycerol (TAG) levels did not change in response to exercise, suggesting that the change in lipid droplet size reflects a switch to a lipid mobilization mode whereby large lipid droplets are broken into smaller ones, and not necessarily an overall lipid depletion. Another hallmark of lipid mobilization in *Drosophila* larvae is the accumulation of lipid droplets in the hepatocyte-like oenocytes [43]. Induction of lipid mobilization from the larval adipose tissue, either by starvation or adipose tissue specific over-expression of the *brummer* lipase, leads to immediate accumulation of lipid droplets in the oenocytes [43]. While the exact mechanism behind this shuttling of lipids is not well characterized, a number of recent studies indicate the pivotal role of the oenocytes in lipid partitioning and systemic regulation of lipid metabolism [44, 45]. Interestingly, larvae subjected to the acute exercise regimen also showed a significant increase in deposition of lipid droplets in the oenocytes, further demonstrating that lipid mobilization is induced in response to acute exercise (Figure 2E-F). It is known that acute exercise may also lead to a metabolic flexibility where nutrient stores are repartitioned from storage sites such as the liver and adipose tissue to help fuel skeletal muscle activity [46]. These physiological changes are known to trigger changes in gene expression that are associated with acute exercise [7]. For instance, acute exercise is known to dramatically increase the expression of interleukin-6 (IL6) in the muscle, one of the very first exercise-induced myokines identified in vertebrates [47] and a known activator of JAK/STAT pathway [48]. There are three cytokines in *Drosophila*: *unpaired* (*upd*) *1*, *upd2* and *upd3*, which are known to activate the JAK/STAT pathway in *Drosophila* [49, 50]. Furthermore, *upd3* has been recently described as a cytokine with homology to interleukin-6 [51]. Hence, we tested if the expression levels of *upd* genes are regulated following acute exercise. We dissected the body wall muscle/cuticle complexes of control and exercised larvae and measured the levels of *upd1*, *upd2* and *upd3* using qRT-PCR and observed a dramatic increase in the expression levels of *upd2* in response to acute exercise (Figure 2G). Expression of *upd3* also increased significantly but was not as pronounced as *upd2*, and the levels of *upd1* remained unchanged (Figure 2G).

Taken together, these observations support the validity of the OMC model to mimic acute exercise-like phenotypes in *Drosophila* larvae.

### Transcriptional responses in the larval muscle in response to acute exercise

Exercise is known to induce robust transcriptional responses within skeletal muscles [7, 52] and proper assessment of such transcriptional changes may depend on the timing or duration after the exercise bouts [53]. To identify gene expression changes that respond immediately to an exercise bout, we chose to assess the transcriptional responses that happen in the body wall muscles in response to our larval acute exercise regimen. Thus, we performed RNAseq on polyA-selected mRNA derived from dissected larval body wall muscles from control and exercised animals at the end of the 2 hours exercise regimen (Figure 3A). Principal component analysis (PCA) of the transcriptome data shows the separation of the control samples from the samples of exercised animals (Figure S3A). The analysis of the differentially expressing genes revealed a total of 2004 genes that changed significantly between the control and experimental samples with adjusted P value < 0.05. Of these, 1190 genes showed a change of 1.5 fold or more, with 672 genes showing an upregulation of 1.5 fold or more and 518 genes showing a downregulation of 1.5 fold or more (Supplementary Table 1, Figure 3B). A heatmap representing these 1190 significantly changing genes show some variability in their expression within the control and experimental replicates (Figure 3B), which may have led to the identification of a lower number of significantly up or downregulated genes than expected. Nevertheless, we did see a significant increase in the expression of both *upd2* and *upd3* as expected from the qRT-PCR results (Figure 2G). To obtain a systems-wide understanding of the biological processes affected by our acute exercise models, we performed a Gene Ontology (GO) enrichment analysis of the genes that showed a statistically significant fold-change of 1.5 or more in expression (Figure 3C). Additionally, we looked at the relationship between the top enriched GO categories and found that these categories primarily mapped to processes such as regulation of growth and cellular component size, response to stimuli or signaling pathways, response to starvation, and morphogenesis (Figure 3D). Interestingly, we identified some significantly enriched GO categories that have physiological relevance to exercise. These categories include processes such as response to wound healing, response to oxidative stress, response to glutathione synthesis, cellular response to unfolded protein, and open tracheal system development (Figure S3B).

As noted above, our exercise regime induced changes in expression of genes that are involved in signaling pathways, growth, and cellular compartment size. These genes most likely represent early responses that lead to physiological changes such as increase in muscle mass, mitochondrial density, and vascularization. Relatedly, the genes that comprise the GO terms related to growth, cell signaling, and tracheal system development might play roles in manifesting these exercise-induced changes. Changes in the expression of genes involved in morphogenesis could be an artifact of the stage at which we exercised the larvae. In holometabolous insects, critical weight is a developmental checkpoint associated with the third instar larvae and determines whether the animal has enough energy reserves to survive metamorphosis [54, 55]. We used mid third instar larvae for the acute exercise experiments. These animals had already cleared the critical weight checkpoint. When starved or stressed, animals that have crossed the checkpoint hasten entry into metamorphosis [55, 56]. Since our animals were exercised in petri dishes that only contained a small volume of 2% sucrose, the animals may have been starved and thus initiated entry into metamorphosis. Although both our control and experimental animals were kept under identical conditions, the additional stress of being contracted could have triggered a stronger starvation response in these animals. Further reducing the duration of the acute exercise paradigm and/or using pre-critical weight larvae might mitigate these changes in metamorphosis genes.

### Analysis of genes that are upregulated by the acute exercise regimen

To determine whether specific biological processes were activated or suppressed in response to exercise, we further separately analyzed genes that were either up- or down-regulated in response to the acute exercise paradigm. Interestingly, GO term enrichment analysis of the 518 downregulated genes yielded very few significantly enriched GO term categories. For example, enrichment analysis of downregulated genes using the widely used GO profiler algorithm identified only 3 broad categories: ion binding; membrane-related genes, and metabolic genes. Similarly, analysis of these genes using another GO term enrichment algorithm, clusterProfiler [57], yielded a single enriched category: drug catabolic process. In contrast, analysis of the up-regulated genes revealed a number of significantly enriched terms. The top 25 highly enriched GO terms show a significant over-representation of genes involved in wound healing, immune response, glutathione metabolic process, and protein folding (Figure 4A). Analysis of the relationships between the various enriched GO terms show that they cluster primarily around processes such as protein folding, amino acid and modified amino acid metabolic processes, and immune response processes (Figure 4B). The top 10 enriched GO categories, including the genes that comprise these groups and the degree of upregulation of each of these genes, is shown in Figure 4C. A similar depiction of genes that comprise an additional six categories that were hand selected for their potential involvement in exercise biology, is shown in Figure S4. These categories include innate immune response, negative regulation of immune response, muscle attachment, tube morphogenesis, response to hypoxia, and negative regulation of proteolysis. Some of these processes, such as enhanced wound healing, are known to benefit from regular exercise [58], and there is increasing evidence suggesting that repeated physical exercise improves immunosurveillance and immunocompetence [59]. Furthermore, one category (“response to hypoxia”) was enriched for several heat shock protein (Hsp) coding genes, which are known to peak after an initial exposure to heat generated as a consequence of exercise in higher vertebrates (see review [60]). Importantly, analysis of the genes that are upregulated revealed several secreted protein coding genes that may represent putative ‘myokines’ (Table 2). Altogether, the gene sets that are activated in response to acute exercise in our OMC model are relevant in the context of human exercise biology.

### Effect of acute exercise on major signaling pathways

Genes involved in cellular signaling pathways are expected to play a key role in mediating both cell autonomous and non-autonomous impacts of acute exercise. Indeed, enrichment of terms such as regulation of response to stimuli and regulation of cell communication in our GO term enrichment analysis indicates that the acute exercise paradigm does impact signaling pathway activities (Figure 3C-D). We analyzed our data to determine the effect of the acute exercise regimen on expression levels of an exhaustive list of genes that are components of the core signaling pathways in *Drosophila*. An enrichment analysis looking for representation of the core signaling pathway components amongst all significantly changing genes (log2FC ≤ −0.6 and ≥ 0.6) show that FGFR signaling and Insulin signaling pathways were most affected (Figure 5A). Similarly, analysis looking for representation of the core signaling pathway components amongst all significantly upregulated genes (log2FC ≥ 0.6) show that JNK, Imd, and TNF-α signaling pathways were strongly affected (Figure 5A). The change in expression of 335 core signaling pathway genes in response to the acute exercise regimen were grouped into 15 core signaling pathways. Of these, 10 pathways had at least 1 gene that showed a significant change of 1.5 fold or more (Figure S5). While many of these genes could be involved in more than one pathway, each gene (with a fold change of 1.5 or more) was represented in only one of the pathways for simplicity in Figure 5B. Importantly, gene sets that are commonly reported as upregulated in healthy or metabolically impaired human subjects in response to aerobic exercise regimens [7] are also over-represented among our acute exercise-induced genes (Figure 5C).

### Concluding Remarks

In summary, we have established an optogenetic muscle contraction (OMC) model in *Drosophila* larvae that provides a fast and effective method for induction of acute exercise regimens. Although exercise models have been developed in adult flies, most of the studies mainly focused on the physiological effects of endurance training. Importantly, studies on the molecular pathways that are activated in muscles upon exercise have not been addressed. Hence, compared to existing exercise models in *Drosophila*, our larval OMC model is unique in 1) genetically inducing exercise in larvae and 2) characterizing muscle-specific transcriptional responses to acute exercise and the molecular pathways that are activated in the exercised muscles. Importantly, the transcriptomic analysis of exercised muscles revealed striking similarities between acute exercise-induced genes in flies and in humans. Altogether, our larval muscle contraction model opens a path for rapid identification and characterization of exercise-induced factors such as myokines.

## ACKNOWLEDGMENTS

We thank the assistance provided by the Microscopy Resources on the North Quad (MicRoN) core and the Sulzberger Columbia Genome Center, Columbia University for RNA sequencing. We thank Drs. Stephanie E. Mohr and Pedro Saavedra for valuable suggestions and comments on the manuscript. We would also like to thank Dr. Rich Binari for assistance in the conduct of this work. This work was supported in part by NIH NIGMS P41 GM132087. N.P. is an investigator of the Howard Hughes Medical Institute.

---

**Figure S1:**
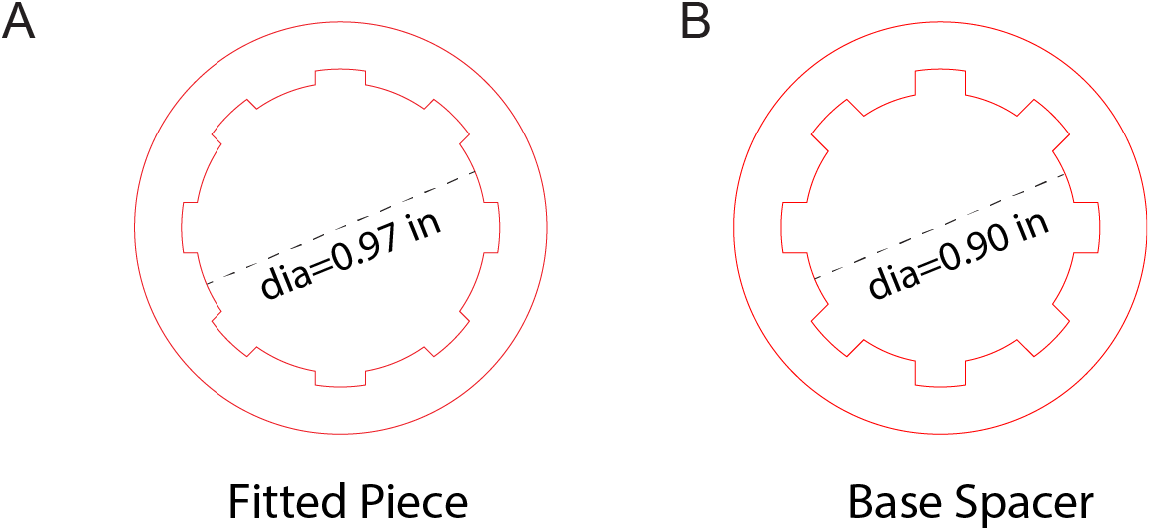
Scaled diagrams for the adapter ring setup. The two rings were stacked such that the cogs of the Fitted piece (A) overlap the troughs in the Base spacer (B) and then joined using acrylic cement to generate the holder. The inner diameter of the rings is 0.97 inches (A) and 0.90 inches (B). Note: The stroke width of the rings has been increased from 0.001 to 0.25 for a better visibility in the manuscript figure. For laser cutting, the lines were actually kept at a dimension of 0.001 inches.

**Figure S2:**
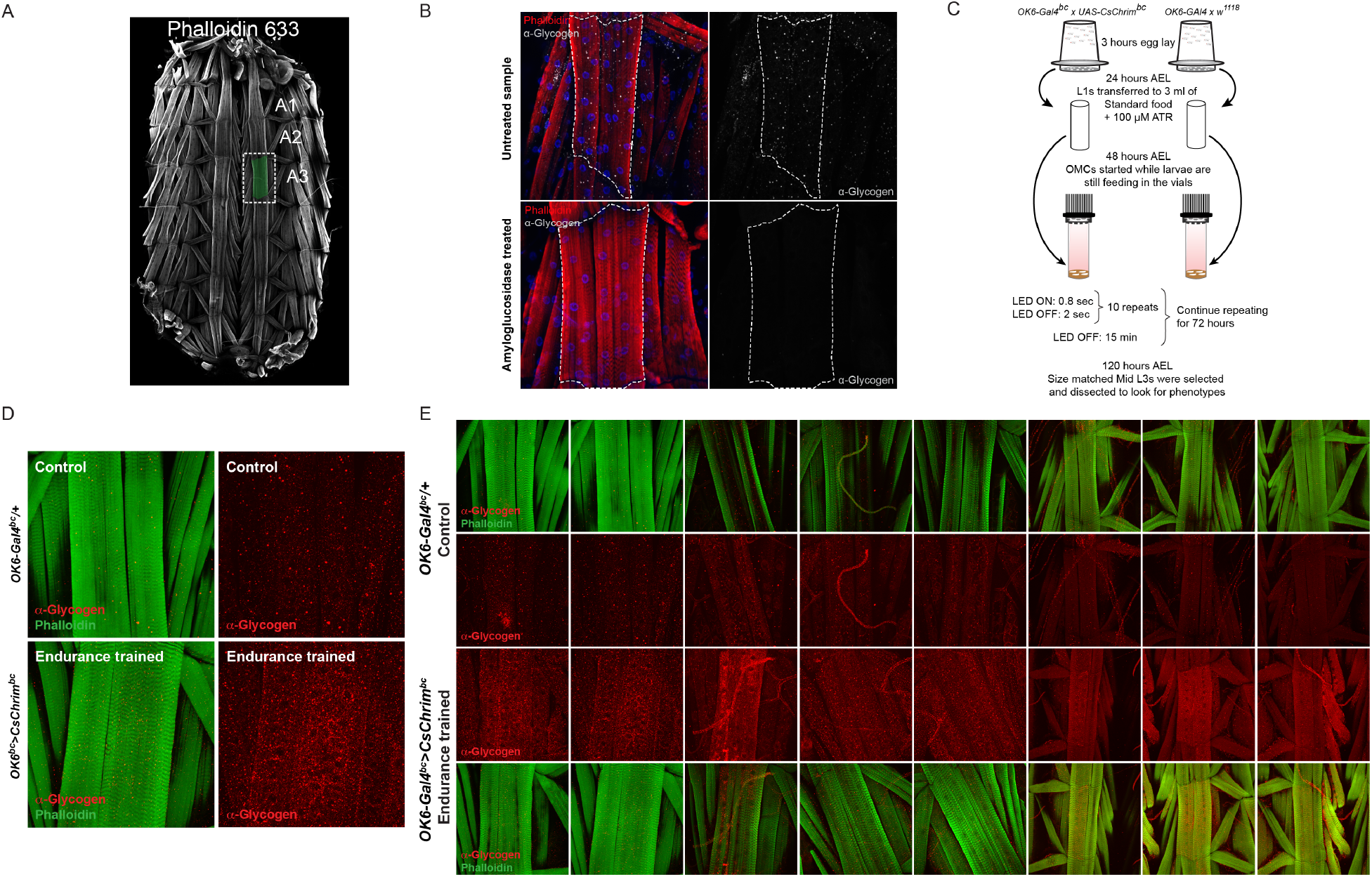
Establishment of an endurance training regimen using OMCs. (A) Representative larval body wall muscle showing the muscle segment A3, which is used for imaging. Body wall muscle is stained with Phalloidin 633 (far red). (B) *w*^*1118*^ larval body wall muscles stained with anti-glycogen antibody (α-Glycogen in grey), phalloidin-594 (red), and DAPI (blue). Sample in the lower panel was treated with amyloglucosidase for 20 min before staining with α-Glycogen antibody. (C) A schematic showing the experimental set up and frequency of OMCs used for modeling of endurance exercise. (D) Anti-Glycogen (α-Glycogen) staining of larval body wall muscles from control animals and animals that were endurance trained. Endurance trained animals show much stronger staining indicating increased glycogen storage in endurance trained muscles. (E) Anti-glycogen (α-Glycogen) staining of control and endurance trained larval body wall muscles. These are biological replicates (n=8) pertaining to (D). Note that the images in the second column are represented in (D).

**Figure S3:**
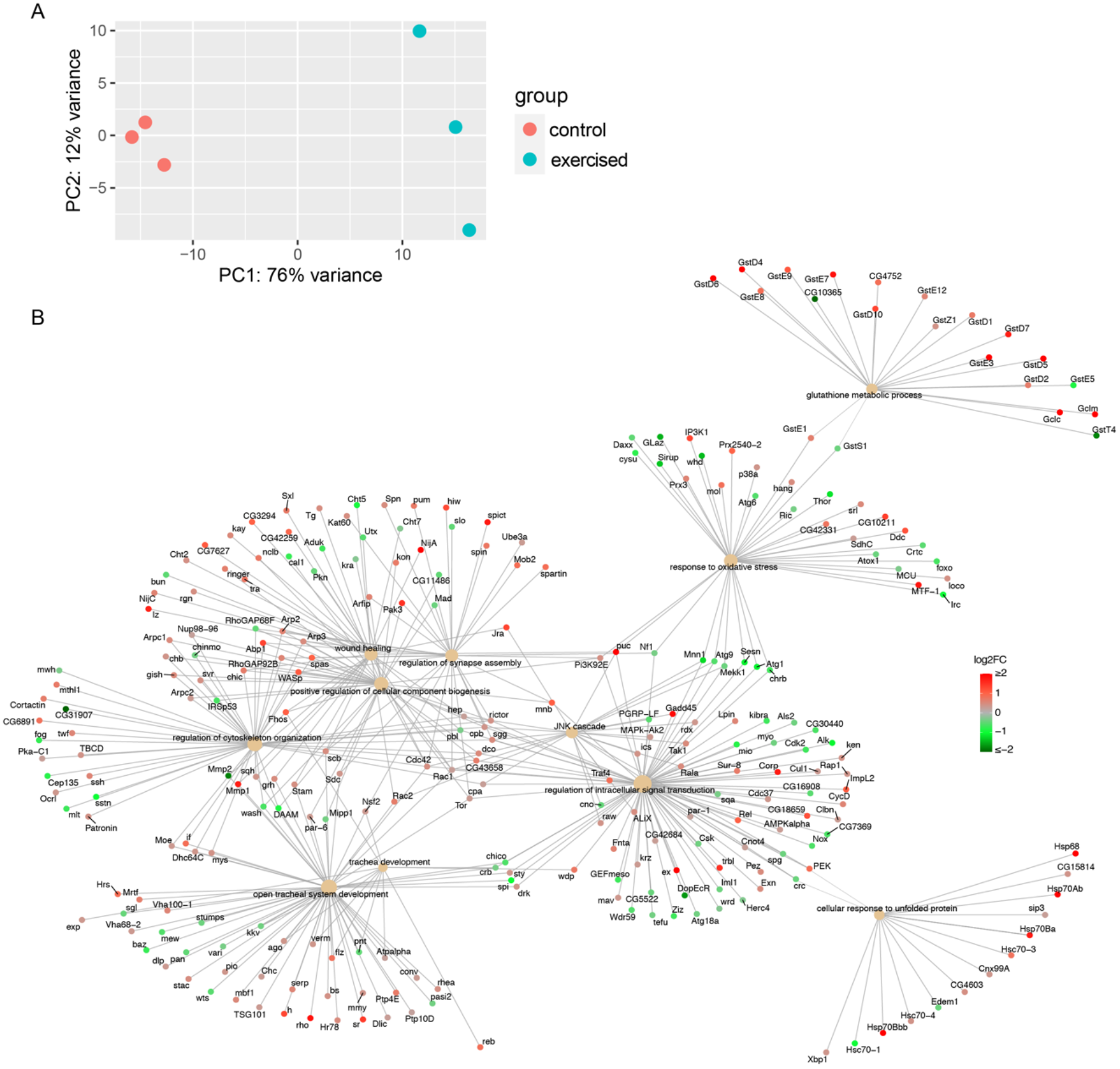
Acute exercise affects genes involved in tracheal development, wound healing, oxidative stress, and glutathione metabolism. (A) Principal Component Analysis (PCA) showing the separation of control and exercised samples using top variance genes. (B) A cnet-plot showing the 10 selected significantly enriched GO terms with the relevant differentially expressing genes. The color of gene node represents the log2 value of fold change with the brightest red representing the log2-FoldChange ≥ 2 and the darkest green representing the log2FoldChange ≤-2

**Figure S4.**
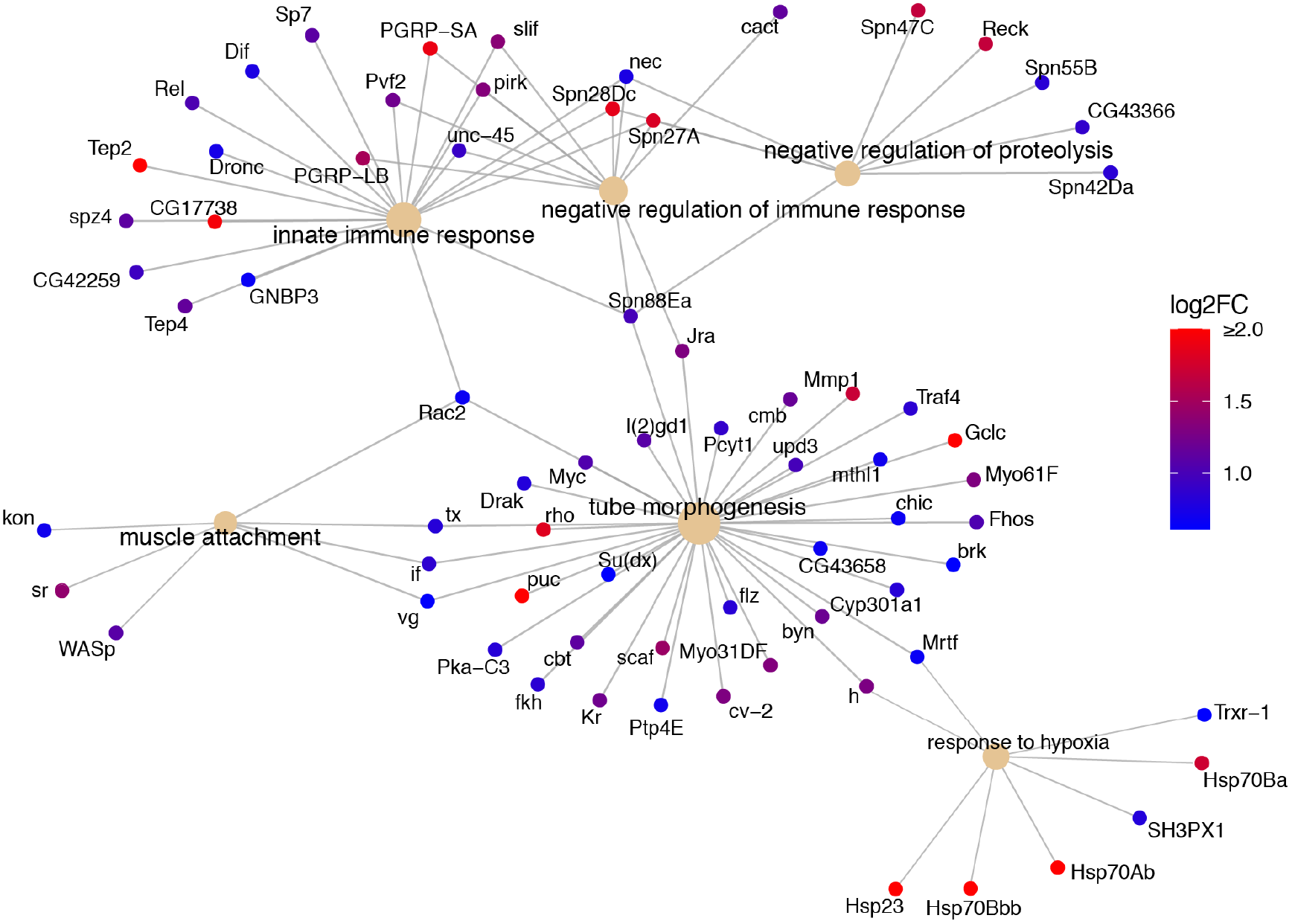
Acute exercise upregulates genes involved in response to hypoxia, tube morphogenesis, and muscle attachment. A cnet-plot showing additional 6 selected GO terms that were significantly enriched for genes that show an upregulation of 1.5 fold or more. The color of gene node represents the log2 value of fold change with the brightest red representing the log2-FoldChange ≥ 2 and the darkest blue representing a log2-FoldChange of 0.60.

**Figure S5:**
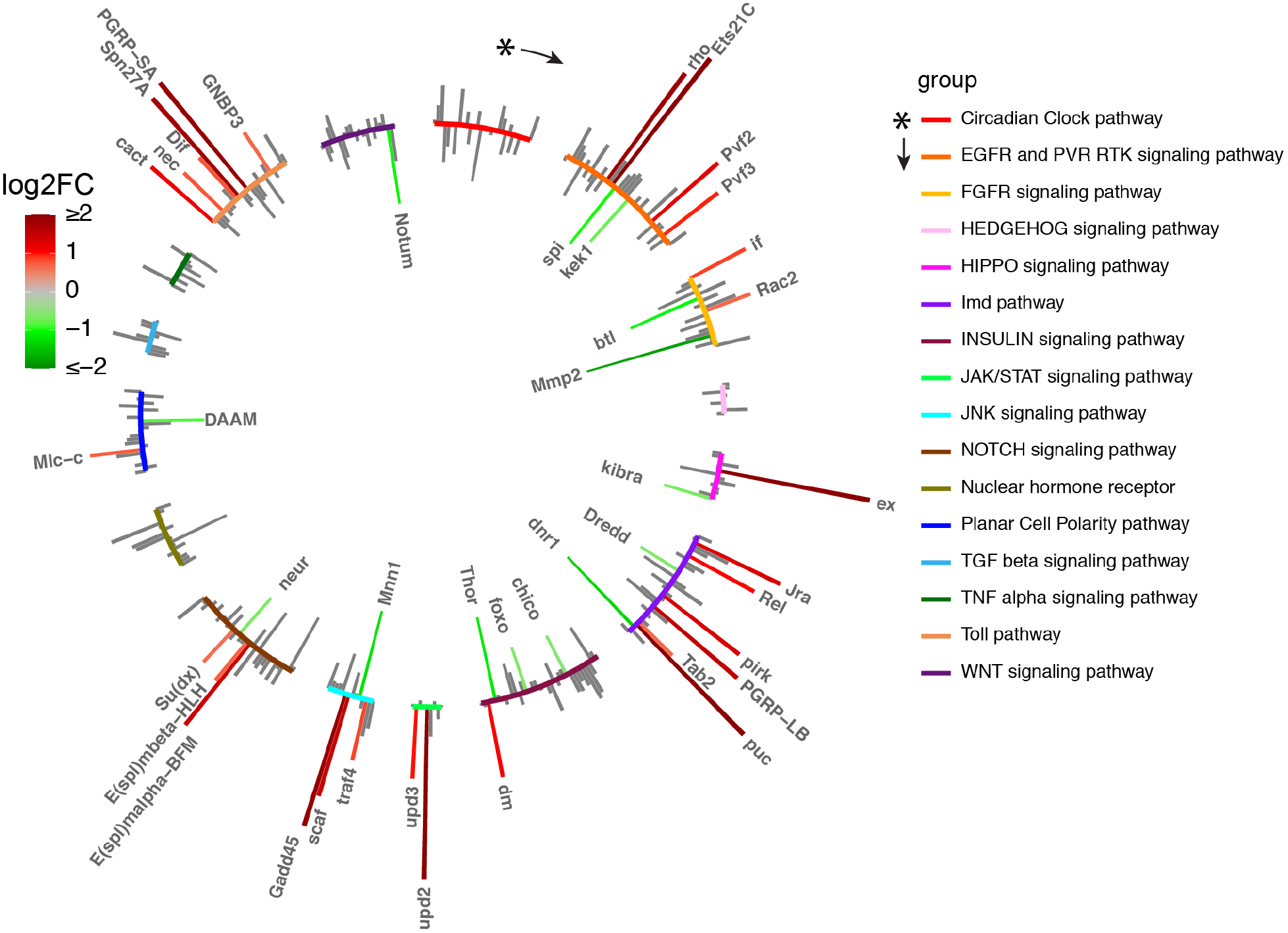
Changes in expression of core signaling pathway genes in response to acute exercise. Genes that show a significant and more than 1.5 fold change in response to exercise are color coded with red showing an increase in expression and green a decrease in expression. Note that some genes are involved in more than one pathway, however, for simplicity of representation these genes have been shown for only one pathway.

## Notes

### Competing Interest Statement

The authors have declared no competing interest.

## References

1. Metter EJ, Talbot LA, Schrager M, Conwit R. Skeletal muscle strength as a predictor of all-cause mortality in healthy men. J Gerontol A Biol Sci Med Sci. 2002;57:B359–365.

2. Nair KS. Aging muscle. Am J Clin Nutr. 2005;81:953–63.

3. Ruiz JR, Ortega FB, Martínez-Gómez D, Labayen I, Moreno LA, De Bourdeaudhuij I, et al. Objectively Measured Physical Activity and Sedentary Time in European Adolescents: The HELENA Study. Am J Epidemiol. 2011;174:173–84.

4. Dempsey PC, Larsen RN, Sethi P, Sacre JW, Straznicky NE, Cohen ND, et al. Benefits for Type 2 Diabetes of Interrupting Prolonged Sitting With Brief Bouts of Light Walking or Simple Resistance Activities. Diabetes Care. 2016;39:964–72.

5. Dempsey PC, Sacre JW, Larsen RN, Straznicky NE, Sethi P, Cohen ND, et al. Interrupting prolonged sitting with brief bouts of light walking or simple resistance activities reduces resting blood pressure and plasma noradrenaline in type 2 diabetes. J Hypertens. 2016;34:2376–82.

6. Gabriel BM, Zierath JR. Circadian rhythms and exercise - re-setting the clock in metabolic disease. Nat Rev Endocrinol. 2019;15:197–206.

7. Pillon NJ, Gabriel BM, Dollet L, Smith JAB, Sardón Puig L, Botella J, et al. Transcriptomic profiling of skeletal muscle adaptations to exercise and inactivity. Nat Commun. 2020;11:470.

8. Krogh-Madsen R, Thyfault JP, Broholm C, Mortensen OH, Olsen RH, Mounier R, et al. A 2-wk reduction of ambulatory activity attenuates peripheral insulin sensitivity. J Appl Physiol Bethesda Md 1985. 2010;108:1034–40.

9. Lanza IR, Short DK, Short KR, Raghavakaimal S, Basu R, Joyner MJ, et al. Endurance exercise as a countermeasure for aging. Diabetes. 2008;57:2933–42.

10. Benatti FB, Pedersen BK. Exercise as an anti-inflammatory therapy for rheumatic diseases-myokine regulation. Nat Rev Rheumatol. 2015;11:86–97.

11. So B, Kim H-J, Kim J, Song W. Exercise-induced myokines in health and metabolic diseases. Integr Med Res. 2014;3:172–9.

12. Saraceni C, Broderick TL. Cardiac and metabolic consequences of aerobic exercise training in experimental diabetes. Curr Diabetes Rev. 2007;3:75–84.

13. Metter EJ, Conwit R, Tobin J, Fozard JL. Age-Associated Loss of Power and Strength in the Upper Extremities in Women and Men. J Gerontol Ser A. 1997;52A:B267–76.

14. Wolin KY, Yan Y, Colditz GA, Lee I-M. Physical activity and colon cancer prevention: a meta-analysis. Br J Cancer. 2009;100:611–6.

15. Monninkhof EM, Elias SG, Vlems FA, van der Tweel I, Schuit AJ, Voskuil DW, et al. Physical activity and breast cancer: a systematic review. Epidemiol Camb Mass. 2007;18:137–57.

16. Borer KT. Physical activity in the prevention and amelioration of osteoporosis in women: interaction of mechanical, hormonal and dietary factors. Sports Med Auckl NZ. 2005;35:779–830.

17. Severinsen MCK, Pedersen BK. Muscle-Organ Crosstalk: The Emerging Roles of Myokines. Endocr Rev. 2020;41:bnaa016.

18. Piccirillo R. Exercise-Induced Myokines With Therapeutic Potential for Muscle Wasting. Front Physiol. 2019;10:287.

19. Jedrychowski MP, Wrann CD, Paulo JA, Gerber KK, Szpyt J, Robinson MM, et al. Detection and Quantitation of Circulating Human Irisin by Tandem Mass Spectrometry. Cell Metab. 2015;22:734–40.

20. Boström P, Wu J, Jedrychowski MP, Korde A, Ye L, Lo JC, et al. A PGC1-α-dependent myokine that drives brown-fat-like development of white fat and thermogenesis. Nature. 2012;481:463–8.

21. Rao RR, Long JZ, White JP, Svensson KJ, Lou J, Lokurkar I, et al. Meteorin-like is a hormone that regulates immune-adipose interactions to increase beige fat thermogenesis. Cell. 2014;157:1279–91.

22. Vinel C, Lukjanenko L, Batut A, Deleruyelle S, Pradère J-P, Le Gonidec S, et al. The exerkine apelin reverses age-associated sarcopenia. Nat Med. 2018;24:1360–71.

23. Hoffmann C, Weigert C. Skeletal Muscle as an Endocrine Organ: The Role of Myokines in Exercise Adaptations. Cold Spring Harb Perspect Med. 2017;7:a029793.

24. Zierath JR, Wallberg-Henriksson H. Looking Ahead Perspective: Where Will the Future of Exercise Biology Take Us? Cell Metab. 2015;22:25–30.

25. Sujkowski A, Wessells R. Using Drosophila to Understand Biochemical and Behavioral Responses to Exercise. Exerc Sport Sci Rev. 2018;46:112–20.

26. Watanabe LP, Riddle NC. New opportunities: Drosophila as a model system for exercise research. J Appl Physiol Bethesda Md 1985. 2019;127:482–90.

27. Damschroder D, Richardson K, Cobb T, Wessells R. The effects of genetic background on exercise performance in Drosophila. Fly (Austin). 2020;14:80–92.

28. Piazza N, Gosangi B, Devilla S, Arking R, Wessells R. Exercise-training in young Drosophila melanogaster reduces age-related decline in mobility and cardiac performance. PloS One. 2009;4:e5886.

29. Tinkerhess MJ, Healy L, Morgan M, Sujkowski A, Matthys E, Zheng L, et al. The Drosophila PGC-1α homolog spargel modulates the physiological effects of endurance exercise. PloS One. 2012;7:e31633.

30. Sujkowski A, Ramesh D, Brockmann A, Wessells R. Octopamine Drives Endurance Exercise Adaptations in Drosophila. Cell Rep. 2017;21:1809–23.

31. Hu Y, Comjean A, Perkins LA, Perrimon N, Mohr SE. GLAD: an Online Database of Gene List Annotation for Drosophila. J Genomics. 2015;3:75–81.

32. Nader GA. Concurrent strength and endurance training: from molecules to man. Med Sci Sports Exerc. 2006;38:1965–70.

33. Nielsen J, Holmberg H-C, Schrøder HD, Saltin B, Ørtenblad N. Human skeletal muscle glycogen utilization in exhaustive exercise: role of subcellular localization and fibre type. J Physiol. 2011;589:2871–85.

34. Baldwin KM, Fitts RH, Booth FW, Winder WW, Holloszy JO. Depletion of muscle and liver glycogen during exercise: Protective effect of training. Pfl1gers Arch Eur J Physiol. 1975;354:203–12.

35. Baba O. [Production of monoclonal antibody that recognizes glycogen and its application for immunohistochemistry]. Kokubyo Gakkai Zasshi. 1993;60:264–87.

36. Zirin J, Nieuwenhuis J, Perrimon N. Role of autophagy in glycogen breakdown and its relevance to chloroquine myopathy. PLoS Biol. 2013;11:e1001708.

37. Luo C-W, Dewey EM, Sudo S, Ewer J, Hsu SY, Honegger H-W, et al. Bursicon, the insect cuticle-hardening hormone, is a heterodimeric cystine knot protein that activates G protein-coupled receptor LGR2. Proc Natl Acad Sci U S A. 2005;102:2820–5.

38. Dewey EM, McNabb SL, Ewer J, Kuo GR, Takanishi CL, Truman JW, et al. Identification of the gene encoding bursicon, an insect neuropeptide responsible for cuticle sclerotization and wing spreading. Curr Biol CB. 2004;14:1208–13.

39. Peabody NC, Diao F, Luan H, Wang H, Dewey EM, Honegger H-W, et al. Bursicon functions within the Drosophila CNS to modulate wing expansion behavior, hormone secretion, and cell death. J Neurosci Off J Soc Neurosci. 2008;28:14379–91.

40. Veverytsa L, Allan DW. Retrograde BMP signaling controls Drosophila behavior through regulation of a peptide hormone battery. Dev Camb Engl. 2011;138:3147–57.

41. Greiwe JS, Hickner RC, Hansen PA, Racette SB, Chen MM, Holloszy JO. Effects of endurance exercise training on muscle glycogen accumulation in humans. J Appl Physiol Bethesda Md 1985. 1999;87:222–6.

42. Hearris MA, Hammond KM, Fell JM, Morton JP. Regulation of Muscle Glycogen Metabolism during Exercise: Implications for Endurance Performance and Training Adaptations. Nutrients. 2018;10:E298.

43. Gutierrez E, Wiggins D, Fielding B, Gould AP. Specialized hepatocyte-like cells regulate Drosophila lipid metabolism. Nature. 2007;445:275–80.

44. Makki R, Cinnamon E, Gould AP. The development and functions of oenocytes. Annu Rev Entomol. 2014;59:405–25.

45. Ghosh AC, Tattikota SG, Liu Y, Comjean A, Hu Y, Barrera V, et al. Drosophila PDGF/VEGF signaling from muscles to hepatocyte-like cells protects against obesity. eLife. 2020;9:e56969.

46. Goodpaster BH, Sparks LM. Metabolic Flexibility in Health and Disease. Cell Metab. 2017;25:1027–36.

47. Pedersen BK, Steensberg A, Schjerling P. Muscle-derived interleukin-6: possible biological effects. J Physiol. 2001;536 Pt 2:329–37.

48. Heinrich PC, Behrmann I, Haan S, Hermanns HM, Müller-Newen G, Schaper F. Principles of interleukin (IL)-6-type cytokine signalling and its regulation. Biochem J. 2003;374:1–20.

49. Wright VM, Vogt KL, Smythe E, Zeidler MP. Differential activities of the Drosophila JAK/STAT pathway ligands Upd, Upd2 and Upd3. Cell Signal. 2011;23:920–7.

50. Harrison DA, McCoon PE, Binari R, Gilman M, Perrimon N. Drosophila unpaired encodes a secreted protein that activates the JAK signaling pathway. Genes Dev. 1998;12:3252–63.

51. Romão D, Muzzopappa M, Barrio L, Milán M. The Upd3 cytokine couples inflammation to maturation defects in Drosophila. Curr Biol. 2021;31:1780–1787.e6.

52. McGee SL, Hargreaves M. Exercise adaptations: molecular mechanisms and potential targets for therapeutic benefit. Nat Rev Endocrinol. 2020;16:495–505.

53. Kuang J, McGinley C, Lee MJ-C, Saner NJ, Garnham A, Bishop DJ. Interpretation of exercise-induced changes in human skeletal muscle mRNA expression depends on the timing of the post-exercise biopsies. 2020.

54. Shingleton AW, Frankino WA, Flatt T, Nijhout HF, Emlen DJ. Size and shape: the developmental regulation of static allometry in insects. BioEssays News Rev Mol Cell Dev Biol. 2007;29:536–48.

55. Mirth CK, Riddiford LM. Size assessment and growth control: how adult size is determined in insects. BioEssays News Rev Mol Cell Dev Biol. 2007;29:344–55.

56. Pan X, Neufeld TP, O’Connor MB. A Tissue- and Temporal-Specific Autophagic Switch Controls Drosophila Pre-metamorphic Nutritional Checkpoints. Curr Biol CB. 2019;29:2840–2851.e4.

57. Yu G, Wang L-G, Han Y, He Q-Y. clusterProfiler: an R package for comparing biological themes among gene clusters. Omics J Integr Biol. 2012;16:284–7.

58. Pence BD, Woods JA. Exercise, Obesity, and Cutaneous Wound Healing: Evidence from Rodent and Human Studies. Adv Wound Care. 2014;3:71–9.

59. Scheffer D da L, Latini A. Exercise-induced immune system response: Anti-inflammatory status on peripheral and central organs. Biochim Biophys Acta Mol Basis Dis. 2020;1866:165823.

60. Hawley JA, Lundby C, Cotter JD, Burke LM. Maximizing Cellular Adaptation to Endurance Exercise in Skeletal Muscle. Cell Metab. 2018;27:962–76.

